# Suppression of *Plasmodium falciparum* infection by *Microsporidia MB* in wild *Anopheles arabiensis* populations

**DOI:** 10.64898/2026.10.16.759541

**Authors:** Thomas Ogao Onchuru, Joseph Gichuhi, Tullu Bukhari, Jack Wardle, Ian Okello, Cassandra Ronnie, Lilian Mbaisi, Priscille Barreaux, Thomas Churcher, Jeremy Keith Herren

## Abstract

Symbiont-based approaches offer a promising strategy for reducing malaria transmission by suppressing *Plasmodium* development in *Anopheles* mosquitoes. *Microsporidia MB*, a naturally occurring microsporidian symbiont of *Anopheles arabiensis*, inhibits *Plasmodium falciparum* development in the laboratory, but its efficacy under natural transmission conditions is unknown. Here we monitored *Microsporidia MB* in wild *An. arabiensis* over 30 months in western Kenya and identified distinct low- and high-intensity infection phenotypes consistent with horizontal and vertical transmission, respectively. Overall, *Microsporidia MB* was associated with approximately 50% lower odds of *P. falciparum* infection. This association strengthened with symbiont intensity: high-intensity infections were associated with approximately 60% lower odds of *Plasmodium* infection compared with approximately 34% for low-intensity infections, identifying symbiont intensity as a key determinant of parasite-blocking efficacy. These findings provide evidence that *Microsporidia MB* suppresses *P. falciparum* infection in wild *Anopheles* populations, providing a key component of the field validation needed to advance *Microsporidia MB* as a candidate for malaria-control applications.

## Introduction

Malaria remains a major impediment to health and economic development across sub-Saharan Africa. In 2024, an estimated 282 million cases were reported globally, resulting in approximately 610,000 deaths, with sub-Saharan Africa accounting for 94% of cases (approximately 265 million) and 95% of deaths (approximately 579,000)[1]. Despite substantial reductions in the malaria burden achieved through the large-scale deployment of insecticide-treated nets (ITNs), indoor residual spraying (IRS), and artemisinin-based combination therapies (ACTs), progress towards malaria elimination has stalled[1,2]. Since 2015, global malaria incidence and mortality have remained consistently high and, in many endemic regions, have increased despite the continued implementation of these interventions[1–3]. This stagnation reflects a convergence of biological and operational challenges, including widespread insecticide resistance in malaria vectors, the emergence and spread of antimalarial drug resistance, climate-driven expansion of vector habitats and transmission seasons, the invasion by *Anopheles stephensi* known to establish in urban areas, and persistent shortfalls in funding for malaria control programmes[1,4–7]. Together, these challenges underscore the significance of expanding the malaria control toolbox with complementary interventions that act through alternative biological mechanisms to further reduce transmission[8].

One promising approach involves leveraging naturally occurring symbiotic microbes to disrupt pathogen development within insect vectors[9]. Symbionts can alter host physiology, immunity, or gut ecology in ways that reduce pathogen transmission, and several have been proposed as candidates for the control of vector-borne diseases[10–12]. The most advanced example is *Wolbachia*, a maternally inherited endosymbiont bacterium that has been introduced into *Aedes* mosquito populations to curb the spread of dengue, Zika, and chikungunya viruses[13,14]. *Wolbachia*-based interventions have demonstrated epidemiological impact in cluster-randomized trials and successful establishment in operational deployments, validating symbiont-based control as a scalable public health strategy[15,16].

The success of *Wolbachia* has established symbiont-based vector control as a viable public health strategy. Although *Wolbachia* infections have been reported repeatedly in wild *Anopheles* populations[17–21], and there is evidence that it may interfere with *Plasmodium*[20,22,23], stably transmitted, high-intensity infections have yet to be achieved in the dominant African malaria vectors (*Anopheles gambiae, Anopheles arabiensis, Anopheles funestus*)[17,21,24], despite some success in establishing transinfected lines in *An. stephensi*[25,26]. This gap has prompted growing interest in other naturally occurring symbionts that might achieve in *Anopheles* what *Wolbachia* has achieved in *Aedes*. *Microsporidia MB*, a symbiont of *An. gambiae s.l.* has emerged as one such promising candidate.

*Microsporidia MB* is a microsporidian, obligate intracellular eukaryotic organism related to fungi, that forms a persistent and non-pathogenic association with *Anopheles* mosquitoes[27]. Unlike *Wolbachia*, *Microsporidia MB* can be transmitted both vertically through maternal inheritance and horizontally via mating[28,29]. Laboratory studies have shown that *Microsporidia MB* substantially reduces *P. falciparum* development in mosquitoes without imposing costs on the hosts[27,30]. These features i.e., robust pathogen suppression, dual transmission modes, and host compatibility, make *Microsporidia MB* a compelling candidate for symbiont-based malaria control. Although *Microsporidia MB* has demonstrated a strong *Plasmodium*-blocking phenotype under experimental conditions, its infection dynamics, transmission, and blocking effects within natural mosquito populations have not been fully characterized. To address this gap, we investigated the prevalence and infection dynamics of *Microsporidia MB* and assessed its association with *Plasmodium falciparum* infection in natural *An. arabiensis* populations. Using entomological surveillance and molecular screening, we assessed whether this naturally occurring symbiont reduces *Plasmodium* infection and development in wild-caught vector mosquitoes. These findings provide insight into the ecological role of *Microsporidia MB* and its potential for development as a sustainable, complementary malaria vector control strategy in sub-Saharan Africa.

## Results

### Distribution, prevalence, and association of *Microsporidia MB* with *Plasmodium* infection in natural *An. arabiensis* populations

A total of 37,879 blood fed and gravid female *Anopheles* mosquitoes collected from the Ahero Irrigation scheme in Western Kenya over a two and a half-year period (January 2022–June 2024) successfully oviposited when they were maintained under laboratory conditions for four days. Using molecular screening of representative mosquitoes from each sampling week, approximately, 96.4% of the mosquitoes collected were identified as *An. arabiensis* (Supplementary Figure 1). The rest of the samples were primarily identified as *An. gambiae s.s* based on COI sequencing (sequences in Supplementary material) while a small fraction remained unresolved due to low-quality sequences. Screening for *Microsporidia MB* revealed consistent presence of the symbiont throughout the study period, with weekly prevalence ranging from 8% to 82% (Figure 1A).

**Figure 1.**
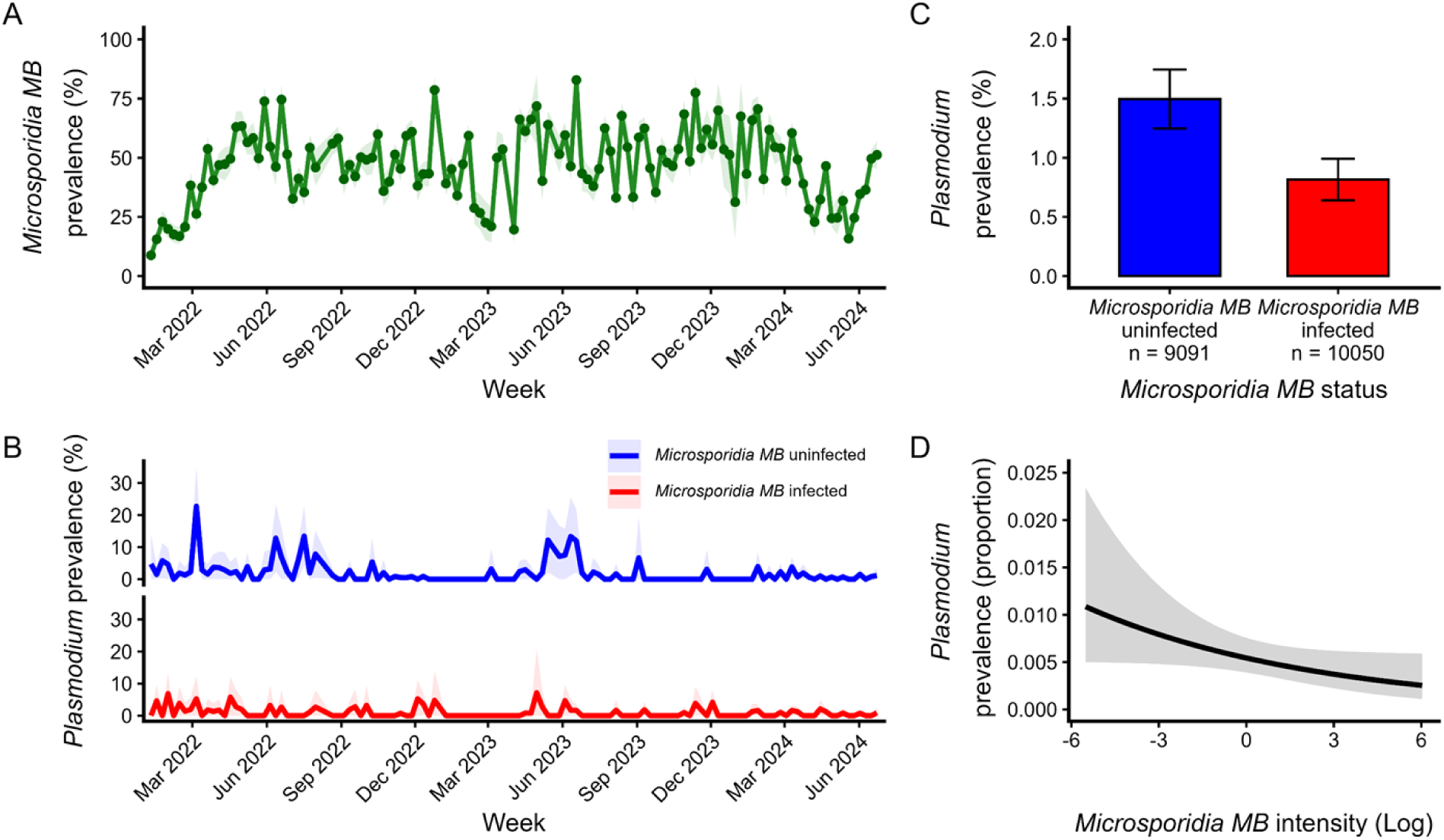
Distribution of *Microsporidia MB* and its association with *Plasmodium falciparum* infection. (A) *Microsporidia MB* endosymbiont is consistently present in *An. arabiensis* populations but with varying prevalence rates. (B) Weekly *Plasmodium* infection prevalence in *Microsporidia MB-*uninfected (blue) and *Microsporidia MB-*infected (red) mosquitoes from January 2022 to June 2024. *Microsporidia MB-*uninfected mosquitoes exhibit higher and more variable *Plasmodium* prevalence, while *Microsporidia MB-*infected mosquitoes have consistently low infection levels throughout the sampling period. (C) Prevalence of *Plasmodium* infection in *An. arabiensis* stratified by natural *Microsporidia MB* infection status. *Microsporidia MB-*infected mosquitoes show 49% lower odds of infection compared to *Microsporidia MB-*uninfected counterparts (OR = 0.51, p < 0.001). (D) Relationship between *Microsporidia MB* intensity and *Plasmodium* infection probability. The black line shows the predicted probability of *Plasmodium* infection from a mixed-effects logistic regression model incorporating continuous *Microsporidia MB* intensity variable and week–year as a random effect. Increasing *Microsporidia MB* infection intensities is associated with reduced odds of infection (OR = 0.881, 95% CI: 0.78–0.995, p = 0.041). Shaded areas represent 95% confidence intervals.

Out of the total number of mosquitoes collected, 9,091 *Microsporidia MB-*uninfected and 10,050 *Microsporidia MB-*infected mosquitoes were screened for *Plasmodium* to determine the association between *Microsporidia MB* presence and infection intensity on *Plasmodium* susceptibility. This analysis only considers *Plasmodium* transmission blocking in the first four days post blood-feeding, when *Plasmodium* oocysts would be expected to be establishing themselves in the midgut of *Anopheles* mosquitoes. Quantifying the complete transmission blocking effect would require detection of sporozoites in the salivary gland 10 days post blood-feeding. Across 124 sampling weeks, *Plasmodium* was detected in 76 weeks (61.3%), while 48 weeks (38.7%) had no detectable infections among screened mosquitoes. Weekly *Plasmodium* prevalence was consistently lower among *Microsporidia MB*-infected mosquitoes (range: 0-7.14%) than among uninfected mosquitoes (range: 0-22%), despite marked temporal variation in both groups (Figure 1B). Overall, *Plasmodium* prevalence was 1.50% (136/9,091; 95% CI: 1.25–1.75%) among *Microsporidia MB*-uninfected mosquitoes compared with 0.82% (82/10,050; 95% CI: 0.64–0.99%) among *Microsporidia MB*-infected mosquitoes (Figure 1C).

To determine parasite species, cytochrome *b* amplicons from a subset of *Plasmodium*-positive samples were subjected to Sanger sequencing. Samples were selected based on sufficient DNA and amplicon quality, while ensuring representation of both *Microsporidia MB*-infected and *Microsporidia MB*-uninfected mosquitoes and periods of relatively high and low *Plasmodium* prevalence. This resulted in 31 samples, comprising 20 of 136 *Microsporidia MB*-uninfected samples and 11 of 82 *Microsporidia MB*-infected samples. Among the sequenced *Microsporidia MB*-uninfected samples, 19 were identified as *P. falciparum* and one as *Plasmodium ovale*. All sequenced *Microsporidia MB*-infected samples were identified as *P. falciparum*. Collectively, *P. falciparum* accounted for 97% (30/31). Sequences to these identifications are available in the Supplementary material. (they will be submitted to NCBI GenBank database and accession numbers provided). All 188 *Plasmodium*-infected mosquitoes that were subjected to host species identification were confirmed as *An. arabiensis*.

The GLMM analysis indicated that *Microsporidia MB*–infected mosquitoes had significantly lower odds of *Plasmodium* infection compared to *Microsporidia MB*–uninfected mosquitoes (OR = 0.507, 95% CI: 0.381–0.671, *p* < 0.001, Figure 1C). When *Microsporidia MB* infection intensity was modelled as a continuous variable, increasing *Microsporidia MB* intensities were significantly associated with lower odds of *Plasmodium* infection (OR = 0.881, 95% CI: 0.78–0.995, *p* = 0.041), after accounting for variation across sampling weeks (Figure 1D). Specifically, each one-unit increase in log-transformed *Microsporidia MB* infection intensity was associated with 11.9% lower odds of *Plasmodium* infection.

### Effect of high and low *Microsporidia MB* infection intensities on *Plasmodium* transmission blocking

We characterized the distribution of *Microsporidia MB* infection intensities in wild-caught *An. arabiensis* vectors and subsequently explored whether variation in infection intensity classes was associated with *Plasmodium* infection. Relative *Microsporidia MB* infection intensities spanned a broad range, exhibiting ratios from <0.1 to 10^6^, with a median infection intensity of 2.2. The overall distribution of *Microsporidia MB* infection intensities in wild-caught *An. arabiensis* was best described by a two-component Gaussian mixture model (GMM; model V, unequal variance) that identified two latent subpopulations (n₁ = 4649; n₂ = 5401) consistent with heterogeneous infection patterns. These clusters were separated by a relative infection intensity cut-off of 0.96 (Figure 2A), classifying individuals into two classes reflecting existence of low- and high-intensity infections. A similar bimodal distribution was observed in colonized mosquitoes, although with a lower cut-off value of 0.25 (Figure 2B). However, *Microsporidia MB* mean infection intensities were also significantly higher in field-caught mosquitoes than in laboratory mosquitoes in both clusters (Welch’s *t*-test: low-intensity cluster, −1.57 in wild caught vs. −1.74 in colonized, *p* < 0.001; high-intensity cluster, 1.84 in wild caught vs. 0.72 in colonized, *p* < 0.001), with a large effect for high-intensity cluster (Cohen’s *d* = 1.09, 95% CI: 0.96–1.23). Interestingly, the two intensity classes in colonized mosquitoes aligned with experimentally controlled transmission modes i.e., vertical or horizontal. Transmission mode strongly predicted *Microsporidia MB* infection intensity, with high-intensity infections predominating in vertically infected mosquitoes (99.1%, 106/107), whereas they occurred in 36.7% (106/289) of horizontally infected mosquitoes. Mosquitoes that acquired *Microsporidia MB* vertically had substantially higher odds of belonging to the high-intensity cluster (Firth logistic regression, OR = 122.2, 95% CI: 33.0–1081.2, *p* < 0.001). Additionally, unestablished low-intensity infections were predominantly associated with horizontally acquired infections and were mostly localized in somatic tissues (Figure 2C, Supplementary Figure 2) whereas high-intensity infections were mainly observed in maternally infected mosquitoes inside germline stem cells and developing oocytes (Figure 2D, Supplementary Figure 2), supporting infection intensity as a useful indicator of the underlying transmission route.

**Figure 2.**
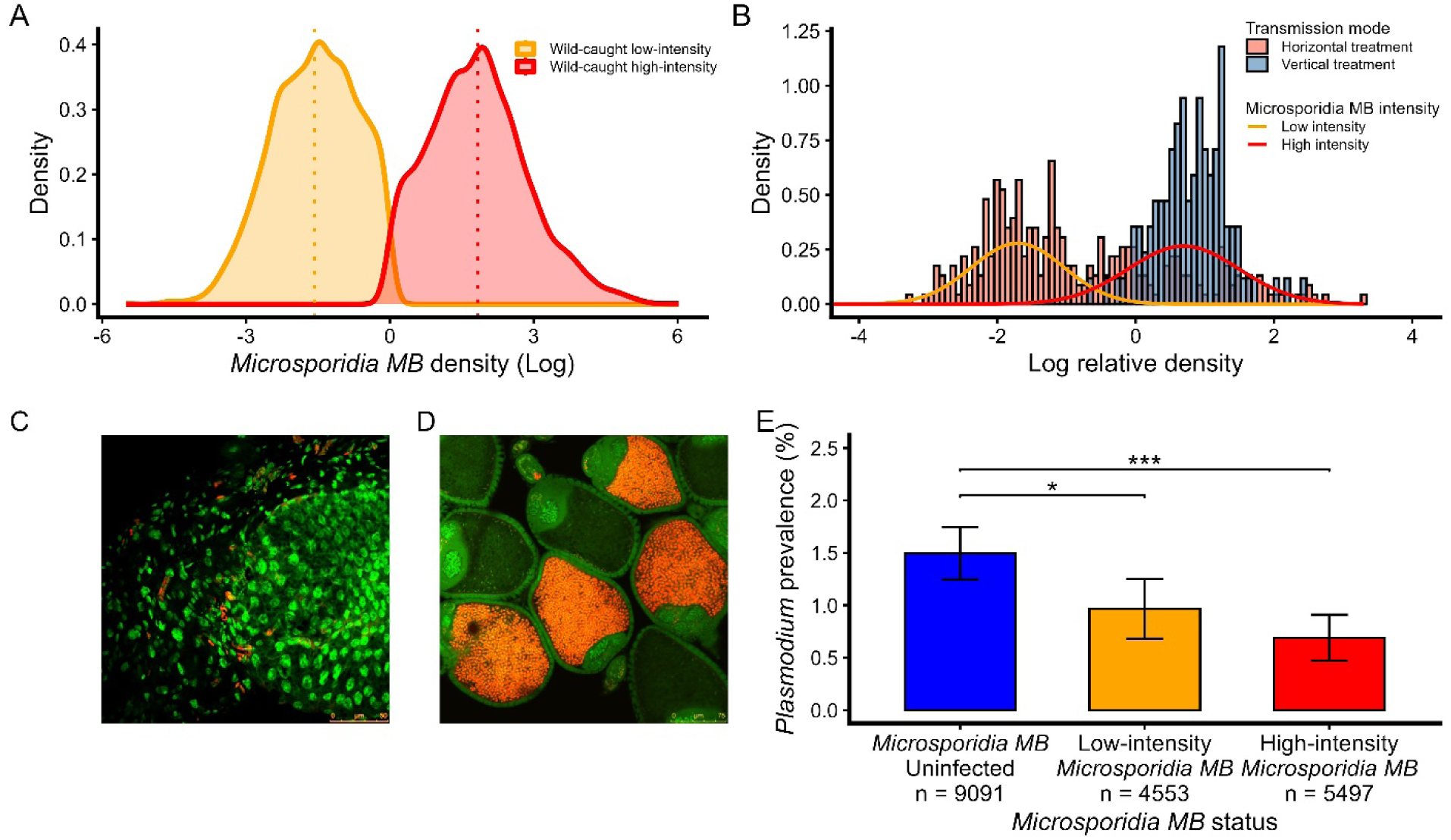
Distribution, transmission mode, localization, and *Plasmodium*-blocking phenotype of *Microsporidia MB* in mosquito populations. (A) Distribution of *Microsporidia MB* infection intensity in wild-caught mosquitoes. Infection intensity distribution was resolved into two populations corresponding to low- and high-intensity MB infections. Dashed vertical lines indicate the mean MB intensity for each population. **(B)** Association between *Microsporidia MB* transmission mode and infection intensity. Mosquitoes from horizontal-transmission treatment are predominantly associated with low-intensity *Microsporidia MB* infections, whereas mosquitoes from vertical-transmission treatment are predominantly associated with high-intensity infections. The proportions of mosquitoes within each intensity class are shown for each transmission mode. **(C-D)** Representative microscopy images of mosquitoes with low-intensity *Microsporidia MB* infection **(C)**, characteristic horizontal transmission and high-intensity *Microsporidia MB* infection **(D)**, characteristic of vertical transmission (red represents *Microsporidia MB* and green represents DNA staining). **(E)** *Plasmodium* prevalence according to *Microsporidia MB* infection status and infection intensity. *Plasmodium* prevalence was significantly lower in *Microsporidia MB* -infected mosquitoes, with the lowest prevalence observed among mosquitoes with high-intensity infections. Bars represent prevalence and error bars indicate 95% confidence intervals; numbers within/below bars indicate the corresponding sample sizes.

To quantitatively evaluate the interaction of infection intensity and localization and define a biologically meaningful threshold, we performed a receiver operating characteristic (ROC) analysis using paired samples with both qPCR-derived *Microsporidia MB* intensity and fluorescence in situ hybridization (FISH) data from the same mosquito (n = 193). *Microsporidia MB* intensity showed excellent discriminatory power for predicting ovarian germline from non-germline infections (AUC = 0.93, 95% CI: 0.89–0.97). Using this method the optimal cut-off (Youden index) was identified as 0.805 with 88% sensitivity and 85% specificity in distinguishing maternal (stem-cell) from horizontal infections (Supplementary Figure 3). Using the quantitatively derived infection cut-off threshold of 0.805, we then evaluated how low and high *Microsporidia MB* infection intensities affect the odds of *Plasmodium* infection from the wild Ahero dataset. The GLMM analysis evaluating the association between *Microsporidia MB* infection intensity and *Plasmodium* infection showed that high-intensity *Microsporidia MB* infections were associated with substantially lower odds of *Plasmodium* infection compared to *Microsporidia MB*-uninfected mosquitoes (OR = 0.403, 95% CI: 0.279–0.582, *P* < 0.001) , while low-intensity *Microsporidia MB* infections were also associated with lower odds of *Plasmodium* infection, although the magnitude of the association was weaker (OR = 0.663, 95% CI: 0.463–0.950, *p* = 0.025) (Figure 2E). Furthermore, high-intensity *Microsporidia MB* infections were associated with significantly lower odds of *Plasmodium* infection than low-intensity MB infections (OR = 0.608, 95% CI: 0.387–0.957, *P* = 0.032), corresponding to approximately 39.2% lower odds, indicating that the strength of the association between *Microsporidia MB* infection and reduced *Plasmodium* infection was greater at higher *Microsporidia MB* infection intensities. As a sensitivity analysis, we repeated the analysis using the wild mosquito population-derived threshold of 0.96 defining low- and high-intensity *Microsporidia MB* infections. The results showed the same overall pattern, with high-intensity infections associated with lower odds of *Plasmodium* infection relative to *Microsporidia MB*-uninfected mosquitoes (OR = 0.376, 95% CI: 0.258–0.550, *P* < 0.001) and low-intensity infections showing a stronger association with lower odds of *Plasmodium* infection (OR = 0.698, 95% CI: 0.491–0.992, *P* = 0.045). In addition, high-intensity infections remained associated with lower odds of *Plasmodium* infection than low-intensity infections (OR = 0.539, 95% CI: 0.342–0.851, *P* = 0.008). Thus, the intensity-dependent association between *Microsporidia MB* infection and *Plasmodium* infection was consistent across the alternative thresholds, indicating that the observed pattern was robust to the choice of threshold. In both analyses, the odds of *Plasmodium* infection were lowest among mosquitoes with high-intensity *Microsporidia MB* infections, intermediate among those with low-intensity infections, and highest among *Microsporidia MB* -uninfected mosquitoes.

### Association of host genetic background with *Microsporidia MB* infection

A total of 20 *An. arabiensis* mosquitoes were included in the host population genetic analysis, comprising 12 *Microsporidia MB*-infected and 8 -uninfected samples. Genome-wide principal component analysis (PCA) revealed substantial overlap between *Microsporidia MB-*infected and uninfected mosquitoes with no distinct clustering according to *Microsporidia MB* infection status (Figure 3A). Similarly, neighbour-joining analyses showed no evidence of *Microsporidia MB*-associated genetic clustering, with *Microsporidia MB-*infected and uninfected individuals interspersed throughout the tree (Figure 3B).

**Figure 3.**
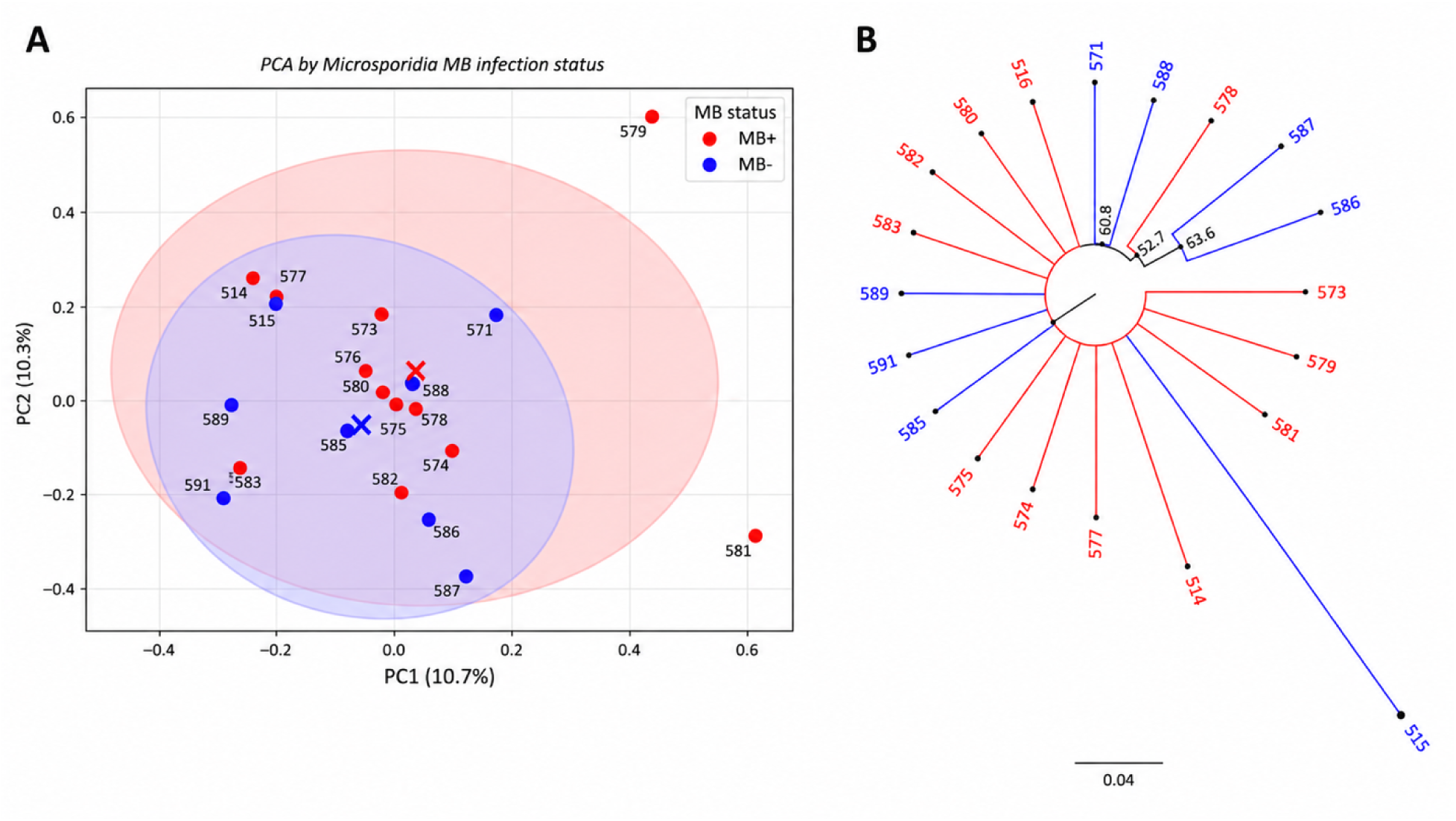
Genome-wide population genetic structure of mosquitoes grouped according to *Microsporidia MB* infection status. **(A)** Principal Component Analysis (PCA) of genome-wide SNP variation. Each point represents an individual mosquito with red and blue indicating *Microsporidia MB-*infected and uninfected individuals, respectively. Ellipses represent the dispersion of samples within each group. The first two principal components explained 10.7% and 10.3% of the total genetic variation, respectively. Extensive overlap between *Microsporidia MB-*infected and uninfected mosquitoes indicates an absence of broad genetic differentiation associated with *Microsporidia MB* infection status. **(B)** Neighbour-joining tree based on genome-wide genetic distances among individual mosquitoes. Tip colours indicate *Microsporidia MB* infection status. *Microsporidia MB-*infected and uninfected individuals are interspersed throughout the tree and do not form distinct clades, providing no evidence of broad population structure associated with *Microsporidia MB*.

Genome-wide genetic differentiation between *MB*-infected and *MB*-uninfected mosquitoes was effectively absent (*F_ST_* ≈ 0). Consistent with this, PERMANOVA analyses detected no significant association between host genetic background and *Microsporidia MB* infection status (pseudo-*F* = 0.96, p = 0.577). Together, these analyses found no evidence of broad genome-wide population structure associated with *Microsporidia MB* infection status in this subset of mosquitoes. Although the limited sample size reduces our ability to detect more subtle genetic differences, we found no indication that *Microsporidia MB* infection was restricted to a major genetically distinct host population. This reduces the likelihood that broad host population stratification explains the observed association between *Microsporidia MB* infection and reduced *Plasmodium* infection.

## Discussion

We investigated whether the *Plasmodium* infection-suppressing phenotype of *Microsporidia MB*, a naturally occurring endosymbiont, is evident in wild-caught *An. arabiensis* mosquitoes from a malaria-endemic region of Kenya. *Microsporidia MB* was consistently present over a period of 2.5 years indicating persistence in natural vector populations. However, its prevalence was heterogeneous ranging from 8% to 82%. This is consistent with what has been reported previously in the area[27,31]. The infection intensities of *Microsporidia MB* in wild-caught *An. arabiensis* spanned a wide range indicating a continuum of newly acquired incipient to well-established high intensity infections. Given the established influence of temperature on *Microsporidia MB* infection dynamics[32], these findings suggest that local environmental conditions may contribute to variable *Microsporidia MB* prevalence and infection intensities observed in field populations.

We also show that *Microsporidia MB* infection is strongly associated with reduced odds of *Plasmodium* infection, with a clear dose-dependent pattern in which high *Microsporidia MB* infection intensities confer substantially greater protection than low infection intensities. Our findings using wild-caught *An. arabiensis* vectors are consistent with previous reports on the ability of *Microsporidia MB* to experimentally block the development and transmission of *Plasmodium* parasites in *An. arabiensis*[27]. The intensity-dependent pathogen blocking phenotype we found mirrors observations reported in other symbiont–vector systems. For instance, in *Wolbachia*–*Aedes aegypti* in vitro and in vivo systems, high intracellular *Wolbachia* titres are essential for achieving robust inhibition of viral replication and dissemination[33–35]. Similarly, differential resistance to viral infections by certain *Wolbachia* variants in *Drosophila melanogaster* is a consequence of their titres[36–39]. Microbiota associated with malaria transmitting mosquitoes can also modulate the anti-*Plasmodium* effects in a dose-dependent manner. A significant decrease of midgut-associated bacteria in *Anopheles gambiae* mosquitoes has been linked with increased levels of *Plasmodium* oocyst infections[12]. Our estimates of parasite inhibition could be conservative because mosquitoes were screened four days post-collection, measuring early oocyst establishment rather than later developmental stages. If *Microsporidia MB* continues to suppress parasite development throughout sporogony, inhibition may increase substantially beyond the four-day time point considered in this study.

We found no evidence of broad genome-wide population structure corresponding to *Microsporidia MB* in this subset of *An. arabiensis.* This is relevant because if *Microsporidia MB* infection were concentrated within a genetically distinct mosquito subpopulation, underlying host population structure could potentially confound the observed association between the symbiont and *Plasmodium* infection. Previous whole-genome studies have shown that broad population structure can be detected in *An. arabiensis* using relatively small sample sets[40]. In our dataset, PCA showed substantial overlap between *Microsporidia MB-*infected and uninfected mosquitoes, while neighbour-joining analysis showed individuals from both groups interspersed throughout the tree. Genome-wide F_ST_ and PERMANOVA similarly showed no evidence of genetic differentiation according to *Microsporidia MB* infection status. Together, these findings provide no evidence that *Microsporidia MB* infection is restricted to a genetically distinct mosquito subpopulation and therefore reduce the likelihood that major host population stratification explains the lower *Plasmodium* prevalence observed among *Microsporidia MB* infected mosquitoes. However, given the relatively small number of mosquitoes included in the genomic analysis, future studies with larger genomic datasets and locus-specific association analyses would therefore be needed to determine whether particular host genetic variants contribute to variation in *Microsporidia MB* infection or its *Plasmodium-*blocking phenotype.

*Microsporidia MB* infection intensities in wild and colonized *An. arabiensis* populations exhibited clear bimodal distributions tightly linked to transmission mode. The cluster with low *Microsporidia MB* infection intensities was consistent with unestablished horizontally acquired *Microsporidia MB* infections while the cluster with high *Microsporidia MB* intensities were strongly enriched in germline-localized infections that were vertically acquired. In the field, despite uncontrolled exposure histories and heterogeneous population structure, the same bimodal pattern emerged, indicating that mixed-mode transmission is robust under natural ecological conditions. The consistent bimodal distribution of *Microsporidia MB* infection intensities in both laboratory and field *An. arabiensis* reflect the symbiont’s mixed transmission ecology where mosquitoes with high and stable *Microsporidia MB* infection intensities that are acquired vertically coexist with those harboring horizontally acquired unestablished and low *Microsporidia MB* infections. Such dual transmission modes are known to generate discrete intensity classes in symbiont systems, especially where vertical transmission ensures high fidelity and amplification across generations, while horizontal acquisition introduces stochasticity in early colonization[41].

The bimodal distribution clusters differed significantly between field and laboratory populations, with field mosquitoes exhibiting substantially higher mean *Microsporidia MB* loads. This divergence may reflect variation in transmission mode, host age, and environmental exposure, which may influence *Microsporidia MB* establishment and proliferation. Field-collected mosquitoes represent a heterogeneous age structure and exposure histories contrast to the relatively synchronized age and controlled exposure conditions of laboratory populations. These findings indicate that natural ecological processes can drive *Microsporidia MB* intensities to levels exceeding those observed under insectary conditions, with important implications for transmission-blocking performance in wild mosquito populations, particularly given our observation that high-intensity infections are more strongly associated with *Plasmodium* suppression. However, because environmental or seasonal parameters were not systematically recorded in this study, we could not directly assess their contributions to variation in *Microsporidia MB* prevalence and intensity or *Plasmodium* susceptibility. Similarly, maternal transmission was assessed only among females that successfully oviposited, potentially limiting our ability to capture transmission dynamics across the full mosquito population. Ongoing longitudinal household-based studies are addressing these questions by simultaneously tracking the interactions between the vector, *Microsporidia MB* and *Plasmodium,* while monitoring vector survival and mortality, environmental parameters and transmission in both male and female mosquitoes. These studies will help clarify how these factors influence *Microsporidia MB* transmission dynamics and the persistence of its *Plasmodium*-suppressing phenotype.

In conclusion, our study provides the first comprehensive field evaluation of the distribution, transmission dynamics and transmission-blocking phenotype of *Microsporidia* MB in natural *Anopheles* populations, demonstrating that naturally infected mosquitoes are substantially less likely to harbour *Plasmodium falciparum*. These findings demonstrate that the parasite-blocking phenotype observed under laboratory conditions is also evident in naturally exposed mosquito populations, supporting its robustness under field transmission conditions. Despite pronounced fluctuations in *Microsporidia MB* prevalence and infection intensities, *Microsporidia MB* was consistently detected in mosquito populations throughout the two-and-a-half-year study period, demonstrating its long-term persistence under natural field conditions. This sustained presence, together with laboratory evidence linking transmission mode with infection intensity, supports the capacity of *Microsporidia MB* to establish and be maintained within mosquito populations over extended periods independent of host population genetic structure. Together, these findings establish *Microsporidia MB* as a naturally occurring, self-sustaining symbiont with strong transmission-blocking activity under real-world ecological conditions, positioning it as a promising strategy with the potential to impact malaria transmission. The observed relationship between *Microsporidia MB* intensity and *Plasmodium* infection highlights the importance of developing strategies that promote the establishment and maintenance of high-intensity *Microsporidia MB* infections in natural vector populations to maximize the potential application of this symbiont as a malaria control tool.

## Material and methods

### Mosquito sampling in Ahero, western Kenya

We conducted a longitudinal entomological survey from January 2022 to June 2024 in villages around the Ahero rice irrigation scheme in western Kenya (0.1740° S, 34.9170° E), a region with perennial malaria transmission and irrigated agriculture that supports year-round mosquito breeding. We collected adult blood-fed female *Anopheles* mosquitoes inside houses by manual aspiration from walls and ceilings between 06:00 and 09:00 hours. Mosquito collections were conducted weekly to capture seasonal variation in mosquito abundance and *Microsporidia MB* and *Plasmodium* infection dynamics. Collections were conducted opportunistically across the study villages, with houses selected at random from a pool of consenting households. On each collection day, a mean of 10 houses (range: 8–12) were sampled, with collections conducted 3–4 days per week. Mosquito collections averaged 306 individuals per sampling week (median = 290; IQR: 191–389) across 124 sampling weeks during the study period.

Mosquitoes were transported alive to the *icipe*-Thomas Odhiambo Campus (Mbita Point) and Duduville Campus (Nairobi) insectaries and maintained under semi-natural conditions in screen houses for four days before they were identified and screened. A four-day holding period was selected to allow complete blood meal digestion and capture established early stage of *Plasmodium* development in the mosquito midgut. *Plasmodium* parasite undergoes the ookinete-to-oocyst transition by approximately 2 days post-infection, with actively growing oocysts evident by 4 days post-infection[42]. Therefore, the four-day interval provided sufficient time for early parasite development and oocyst establishment while providing sufficient time to process large numbers of field-collected mosquitoes with minimal mortality under semi-natural conditions. Mosquitoes were morphologically identified using established morphological keys[43], with species identity independently confirmed by species-specific PCR assays[44] in approximately five randomly selected mosquitoes per week throughout the collection period. For subsequent assessment of the association between *Microsporidia MB* and *Plasmodium* infection, only female mosquitoes that successfully oviposited within four days of collection were included in the analysis. Females that did not oviposit within this period were excluded

### *Microsporidia MB* prevalence and infection intensities

DNA was extracted from individual mosquitoes using the protein precipitation method (Supplementary material). The extracted DNA was used to determine *Microsporidia MB* infection status in all samples using a quantitative PCR (qPCR) assay targeting *Microsporidia MB 18S* rRNA gene region using specific primers (MB18SF: CGCCGGCCGTGAAAAATTTA and MB18SR: CCTTGGACGTGGGAGCTATC) as previously described[27]. The proportion of *Microsporidia MB-*infected mosquitoes per collection week was used to assess prevalence. Across the 124 sampling weeks, an average of 61.2% of *Microsporidia MB*-infected mosquitoes were subsequently assessed for infection intensity (median, 60.9%; IQR, 41.6–80.0%). The mosquito *ribosomal protein S7* gene (primers, S7F: TCCTGGAGCTGGAGATGAAC and S7R: GACGGGTCTGTACCTTCTGG) was amplified by qPCR and used to normalize *Microsporidia MB* DNA quantities relative to host DNA to determine infection intensity. The qPCR reaction set up and conditions for the detection and quantification of *Microsporidia MB* infection intensities are outlined in supplementary material.

### *P. falciparum* prevalence in *Microsporidia MB* infected and uninfected mosquitoes

All *Microsporidia MB*–infected mosquitoes whose infection intensities were determined and an approximately equal number of *Microsporidia MB*–uninfected mosquitoes per collection day were screened for *Plasmodium* infection. This allowed for the determination of the effect of *Microsporidia MB* intensities on *Plasmodium* prevalence relative to uninfected mosquitoes. A qPCR reaction targeting the *Plasmodium* mitochondrial cytochrome *b* gene was used for assessing *Plasmodium* infection[27].

To determine the identity of *Plasmodium* species, Sanger sequencing of the cytochrome *b* amplicons from both *Microsporidia MB*-infected and uninfected mosquitoes was conducted. Sequence-confirmed samples were then used as species-specific positive controls to validate qPCR assays for detecting infections and identifying *Plasmodium* species in all remaining mosquito samples (Supplementary material). The prevalence of *Plasmodium* infections in *Microsporidia MB-*infected and uninfected groups was compared to determine the effect of the symbiont on *Plasmodium*.

### Defining *Microsporidia MB* infection intensity classes and their biological thresholds

To investigate whether the effect of *Microsporidia MB* on *Plasmodium* infection varies with symbiont intensity, we first characterized the distribution of *Microsporidia MB* infection intensities in field-collected and laboratory-reared *An. arabiensis*. Infection intensity was quantified by qPCR and analyzed using Gaussian mixture models (GMMs) to identify latent subpopulations corresponding to distinct low and high infection intensity classes. Field-collected mosquitoes were used to determine whether intensity classes occurred in natural populations. In parallel, laboratory populations were used to determine whether similar intensity classes occurred under controlled conditions and whether they were associated with the mode of symbiont acquisition. For these analyses, mosquitoes were generated in which *Microsporidia MB* was acquired exclusively through either horizontal (male-to-female) or vertical (maternal) transmission. Horizontally infected female treatment was produced by separating female pupae from a *Microsporidia MB*-free colony, maintaining them as virgins, and exposing 2-day-old adults to 5-day-old infected males for seven days. Vertically infected treatment was obtained from an infected colony by separating male and female pupae and maintaining adults on 6% glucose. Infection intensities from these laboratory populations were incorporated into the GMM analyses to evaluate whether transmission route contributed to the observed intensity structure and whether the identified subpopulations corresponded to horizontally and vertically acquired infections. Additionally, fluorescence in situ hybridization (FISH) protocol (Supplementary material) was used to localize *Microsporidia MB* infections following vertical and horizontal transmission[28].

To establish a biologically meaningful threshold separating low- and high-intensity *Microsporidia MB* infections, we investigated whether variation in infection intensity was associated with symbiont transmission mode. Tissue localization was used as a proxy for transmission route, whereby germline-localized infections were considered indicative of vertical transmission and non-germline-localized infections indicative of recent horizontal acquisition. Female mosquitoes from a colony containing mixed-mode infections were dissected and the paired ovaries separated. To control for inter-individual variation in infection intensity, one ovary from each mosquito was used for qPCR quantification of *Microsporidia MB* intensity, whereas the paired ovary was analyzed by fluorescence in situ hybridization (FISH) to determine infection localization. Because *Microsporidia MB* intensity is generally uniform between paired ovaries, localization patterns obtained by FISH could be directly linked to the corresponding qPCR-derived intensity measurements from the paired ovary.

These localization-defined categories were then used as biological labels in receiver operating characteristic (ROC) analysis to evaluate the ability of *Microsporidia MB* intensity to discriminate between germline- and non-germline-localized infections. The optimal threshold was identified using the Youden index, maximizing sensitivity and specificity. This ROC-derived cutoff was subsequently used to classify field-collected mosquitoes into low- and high-intensity infection groups for downstream analyses of *Plasmodium* infection.

### Influence of host genetic background on *Microsporidia MB* infection

To determine whether host genetic background contributed to variation in *Microsporidia MB* infection and thereby influencing *Plasmodium* susceptibility, we compared genome-wide host genetic variation among 20 *An. arabiensis* mosquitoes grouped according to *Microsporidia MB* infection status (12 MB+, 8MB-). Host species identity was confirmed prior to whole-genome sequencing during sample selection using morphological identification and species-specific PCR and was subsequently verified using genome-wide SNP data following whole-genome sequencing. Only *An. arabiensis* individuals were included in downstream analyses. Population genetic structure was assessed using a randomly selected subset of 100,000 high-quality biallelic single nucleotide polymorphisms (SNPs) from the genome-wide variant dataset. Principal component analysis (PCA), neighbour-joining clustering based on pairwise genetic distances, genome-wide fixation index (*F_ST_*) estimates, and permutational multivariate analysis of variance (PERMANOVA) were used to evaluate genetic differentiation among infection groups.

Clustering patterns, genetic distances, and measures of population differentiation were examined to determine whether *Microsporidia MB* infection status was associated with broad host population structure. This analysis was used to assess whether *Microsporidia MB* was concentrated within a genetically distinct mosquito background, which could confound interpretation of the observed association between the symbiont and reduced *Plasmodium* infection.

### Data analysis

The distribution of *Microsporidia MB* infection intensities in field and colonized *An. arabiensis* populations was assessed using a Gaussian Mixture Model (GMM), which identified latent subpopulations within natural vector population. qPCR-derived relative infection intensities were fitted to models containing one to five Gaussian components using the mclust package in R (version 4.0.0). Model selection was guided by the Bayesian Information Criterion (BIC), with the best-fitting model corresponding to the lowest BIC value. Individual mosquitoes were classified into infection categories based on posterior probabilities from the fitted model. This approach was used to identify natural clusters of low versus high *Microsporidia MB* infection intensities, reflecting heterogeneity in infection patterns within natural populations. The association between *Microsporidia MB* infection intensity and transmission mode was evaluated using Firth penalized logistic regression, with infection intensity (low versus high) as the outcome and transmission mode (horizontal versus vertical) as the predictor. Effect sizes were reported as odds ratios (ORs) with 95% confidence intervals (CIs).

We used mixed-effects logistic regression models (GLMMs) to examine the association between *Microsporidia MB* infection and *Plasmodium* presence. First, we assessed whether the presence of *Microsporidia MB* affected the probability of *Plasmodium* infection, as defined by PCR positivity 4 days following collection. We included sampling week-year as a random effect (the same week from different years are considered independent) to account for temporal variation in mosquito collections. Second, we investigated whether *Microsporidia MB* infection intensity was associated with *Plasmodium* prevalence by including *Microsporidia MB* intensity as a continual variable and then into three different intensity groups (as a categorical variable, be-it uninfected, low or high intensity) with sampling week-year as a random effect.

## Ethical consideration

Ethical approval was granted from Kenya Medical Research Institute’s Scientific and Ethics Review Unit to collect mosquitoes from houses following verbal consent (NON-KEMRI 4958).

## Data availability

All data supporting the findings of this study are available within the article, its Supplementary information file, and Figshare data repository with the identifier https://doi.org/10.6084/m9.figshare.33593668

## Acknowledgements

The authors would like to acknowledge Aclaine Shisia, Erick Odoyo, David Alila, Charles Amara, Vincent Gadi, Robinson Kisero, Gerald Ronoh, Edward Makhulu, Fidel Gabriel Otieno, Stancy Ombonya, Mwatum Alakonya for assisting with fieldwork and sample processing, Jeniffer Thiongo and Peris Waweru for assisting with mosquito rearing, Faith Kyengo and Rose Marubu for administrative support. We would also like to thank the Sinkins lab group members (University of Glasgow) for discussions about *Microsporidia MB*.

## Author Contributions

TOO: conceptualization, molecular assays, data analysis, writing the initial draft, revising, and funding acquisition

JG: Conceptualization, data validation and analysis

TB: Conceptualization, analysis, writing and reviewing, and supervision

JW: data analysis and reviewing

IO: molecular assays and validation

CR: molecular assays and validation

LM: data analysis, writing, and reviewing

PB: data analysis and reviewing

TC: data validation and analysis, reviewing, supervision

JKH: Conceptualization, analysis, validation, writing and reviewing, supervision, and funding Acquisition

